# Determinants of the microbiome spatial variability in chronic rhinosinusitis

**DOI:** 10.1101/2022.10.02.509831

**Authors:** Joanna Szaleniec, Valentyn Bezshapkin, Agnieszka Krawczyk, Katarzyna Kopera, Barbara Zapała, Tomasz Gosiewski, Tomasz Kosciolek

## Abstract

**Background:** The sinus microbiome in patients with chronic rhinosinusitis (CRS) is considered homogenous across the sinonasal cavity. The middle nasal meatus is the recommended sampling site for 16S rRNA sequencing. However, individuals with unusually high between-site variability between the middle meatus and the sinuses were identified in previous studies. This study aimed to identify which factors determine increased microbial heterogeneity between sampling sites in the sinuses.

**Methodology:** In this cross-sectional study samples for 16S rRNA sequencing were obtained from the middle meatus, the maxillary and the frontal sinus in 50 patients with CRS. The microbiome diversity between sampling sites was analysed in relation to the size of the sinus ostia and clinical metadata.

**Results:** In approximately 15% of study participants, the differences between sampling sites within one patient were greater than between the patient and other individuals. Contrary to a popular hypothesis, obstruction of the sinus ostium resulted in decreased dissimilarity between the sinus and the middle meatus. The dissimilarity between the sampling sites was patient-specific: greater between-sinus differences were associated with greater meatus-sinus differences, regardless of the drainage pathway patency. Decreased spatial variability was observed in patients with nasal polyps and extensive mucosal changes in the sinuses.

**Conclusions:** Sampling from the middle meatus is not universally representative of the sinus microbiome. The differences between sites cannot be predicted from the patency of communication pathways between them.

## Introduction

The significance of the microbiome for the development of chronic rhinosinusitis (CRS) is unclear. CRS is a complex inflammatory disease and treatment outcomes are influenced by multiple factors (1, 2), while bacteria are believed to cause exacerbations and contribute to the recalcitrance of CRS (1, 3, 4). The effectiveness of antimicrobial treatment and reliability of research depends on representative sampling of the sinonasal microbiome.

It is generally assumed that microbiome samples from the middle nasal meatus are representative of the sinuses (1, 5). In CRS, however, the communication between the sinuses and the middle meatus is often impaired. Previously, we showed that the results of the middle meatus culture in patients with CRS were discordant with maxillary sinus culture in 20% of cases and frontal sinus culture in 34% of cases (6).

In most studies on the sinonasal microbiome that were conducted using 16S rRNA sequencing, the authors observed only small or insignificant differences between the sinuses and the middle meatus (7–10). Nevertheless, several researchers noted larger dissimilarity between sampling sites in some patients (7, 11). The causes of increased between-site differences in certain individuals have not been studied before.

This study aimed to identify the factors that determine the heterogeneity of the sinonasal microbiome. Samples for 16S rRNA sequencing were obtained from the middle meatus, the maxillary sinus, and the frontal sinus. To evaluate the impact of the anatomical separation of the subsites, we compared three groups of patients: (a) with narrow sinus ostia, (b) with blocked sinus ostia, and (c) with wide sinus ostia after previous surgery. Subsequently, we evaluated the relationship between other clinical metadata and microbiome diversity. Obstruction of the sinus ostia did not result in increased differences between the sinuses and the middle meatus. Decreased between-site microbiome variability was associated with certain clinical characteristics of the patients (nasal polyps, extensive opacification of the sinuses).

## Materials and methods

### Sample collection

In this cross-sectional observational study, samples were collected from patients with CRS during endoscopic sinus surgery between October 2018 and June 2019 at the University Hospital in Krakow. CRS was defined according to the EPOS 2012 guidelines (12). All of the participants had not improved after medical treatment and were scheduled for surgery. Patients who received antibiotics one month before the surgery were excluded. Clinical data collected for the patients included: age, time since CRS onset, nasal polyps, comorbidities (asthma, aspirin-exacerbated respiratory disease, gastroesophageal reflux, allergy), recent steroid use, history of recurrent exacerbations and previous sinus surgeries, radiological staging (Lund-Mackay score (13)) and self-evaluation of the symptoms on the visual analogue scale.

The swabs were collected under endoscopic guidance from 3 sites (the middle nasal meatus, the maxillary sinus and the frontal sinus on the same side) in 50 patients which provided a total of 150 samples. If the sinus was blocked, the swab was collected immediately after the surgical opening of its ostium. Contact with the nasal vestibule or other sites was strictly avoided. The samples were transported on ice to the laboratory where they were stored at −80oC. The study was approved by the Jagiellonian University Bioethics Committee (1072.6120.78.2018, 20.04.2018).

### DNA isolation, library preparation and sequencing

The swabs were thawed and vigorously shaken in 1 mL of saline. Afterwards, the samples were treated with lysozyme (1□mg/mL) and lysostaphin (0.1□mg/mL) enzymes (Sigma-Aldrich, Poznan, Poland) at 37□°C for 20□min to digest the bacterial cell walls. Further, the samples were subjected to DNA extraction using a Mini Genomic DNA isolation kit (A&A Biotechnology, Gdynia, Poland). The concentration and purity of DNA isolates were determined spectrophotometrically for A260 nm and A260nm / 280nm ratio using NanoDrop (Thermofisher, Waltham, MA USA).

Libraries were prepared strictly according to Illumina’s protocol (San Diego, CA, USA) and Kowalska-Duplaga *et al*. (14, 15).

Primers (Genomed, Warsaw, Poland) specific to the V3 and V4 16S rRNA sequences of bacteria were used:

(F) 5′TCGTCGGCAGCGTCAGATGTGTATAAGAGACAGCCTACGGGNGGCWGCAG3′ (R)5′

GTCTCGTGGGCTCGGAGATGTGTATAAGAGACAGGACTACHVGGGTATCTAATCC 3′

After purification and concentration measurement, libraries were pooled and sequenced using the MiSeq sequencer (Illumina, San Diego, CA, USA).

In one sample from the middle meatus, amplifying the library was impossible due to too low DNA concentration - it was excluded from further analysis.

### Data and statistical analysis

The sequencing results were processed using a pipeline available within QIIME 2 (16). Truncation was performed at 290 bp length for forward reads and at 250 bp length for reverse reads to avoid the technical quality drop with the increased base position. Paired-end sequences were denoised using DADA2 via q2-dada2 and merged with minimal overlap of 12 base pairs. The phylogenetic tree was constructed using q2-fragment-insertion with the SEPP reference database based on SILVA 128. After analysis of rarefaction curves, the sampling depth of 20,504 reads per sample was chosen. The rarefaction procedure resulted in the exclusion of additional 14 samples from the analysis (7 from the middle meatus, 5 from the maxillary sinus, and 2 from the frontal sinus).

The measure of microbiome diversity within each sample (alpha diversity) used in this study was Faith’s phylogenetic diversity which incorporates phylogenetic differences between the taxa identified in the sample.

The disparities in microbiome composition between pairs of samples (beta diversity) were measured using two dissimilarity metrics: Bray-Curtis and weighted UniFrac (wUniFrac). Bray-Curtis dissimilarity quantifies compositional differences between biological communities based on counts at each site. The values of the metrics range from 0 to 1, where 0 indicates that the samples are identical and 1 means that the two samples do not share any species. The wUniFrac distance additionally incorporates information on the phylogenetic distances between organisms. The lowest wUniFrac value is 0 if the communities do not differ. Larger values indicate greater differences and the maximal value may exceed 1 (17).

Phylogenetic Investigation of Communities by Reconstruction of Unobserved States (PICRUSt2) was used for the prediction of the functional potential of the microbial communities. This computational tool allows for the prediction of metagenomic functional profiles from 16 rRNA sequencing data via hidden state prediction on a constructed phylogenetic tree. Although prediction accuracy is varied depending on the microbial environment, PICRUSt2 was shown to be highly efficient within the human organism (18).

The normality of distribution was assessed using the Shapiro-Wilk test. The t-Student test and Pearson correlation were used for biodiversity measures that had a normal distribution. Wilcoxon signed-rank test, Mann-Whitney-Wilcoxon test, Kruskal-Wallis test and Spearman’s rank correlation were used for biodiversity measures that did not have a normal distribution.

The differences in microbiome composition between subgroups of samples were further explored using permutational multivariate analysis of variance (PERMANOVA) using the adonis function.

Data is available at the European Nucleotide Archive (ENA): https://www.ebi.ac.uk/ena/browser/view/PRJEB55924

The Strengthening The Organization and Reporting of Microbiome Studies (STORMS) Checklist can be found at https://zenodo.org/record/7092029#.YzF6CuxBx6o

The code is available at: https://github.com/bioinf-mcb/spatial_sinus_microbiome

## Results

### Clinical characteristics of the study group

Fifty patients with CRS were enrolled in the study. Samples were collected during endoscopic sinus surgery from the middle meatus, maxillary sinus and frontal sinus. The study participants’ demographics and clinical characteristics are shown in Table 1.

**Table 1.** Demographic and clinical characteristics of the 50 study participants.

|  |  |
| --- | --- |
| Age | 19-83 (mean 49) |
| Gender | female: 25, male: 25 |
| Previous endoscopic sinus surgery | 24 |
| CRS with nasal polyps | 21 |
| Computed tomography – Lund-Mackay score (0-24, greater values indicate greater extension of the sinus opacification) | 2-24 (mean 13) |
| Comorbidities | asthma: 20<br>aspirin-exacerbated<br>respiratory disease: 9<br>allergy: 24<br>gastroesophageal reflux: 15 |

### Variability between sampling sites

In every patient, the Bray-Curtis and weighted UniFrac distances between the following pairs of samples were calculated:

a. middle meatus-maxillary sinus distance,
b. middle meatus-frontal sinus distance,
c. maxillary sinus-frontal sinus distance.

The distribution of the beta-diversity indexes was not normal. Therefore, nonparametric tests were used in the analysis. The values of the weighted UniFrac and Bray-Curtis dissimilarity are shown in Table 2. Although the median values of dissimilarity measures were 0.28-0.34 for the weighted UniFrac and 0.22-0.26 for the Bray-Curtis metrics, in some patients we observed much higher values of the indexes. For example, the maximal values of the Bray-Curtis metric reached 0.75 for the middle meatus-maxillary sinus distance and 0.92 for the middle meatus-frontal sinus.

**Table 2.**
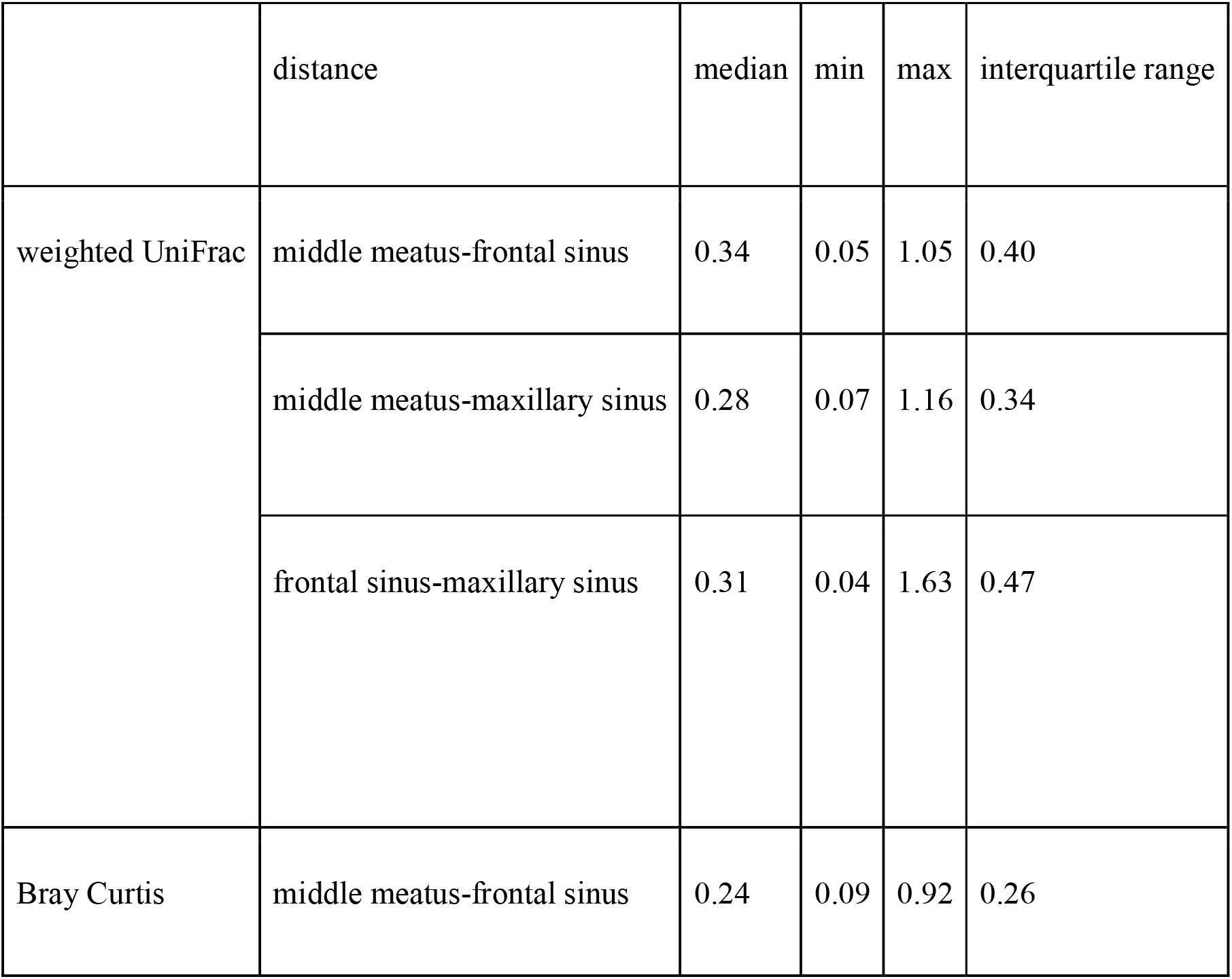
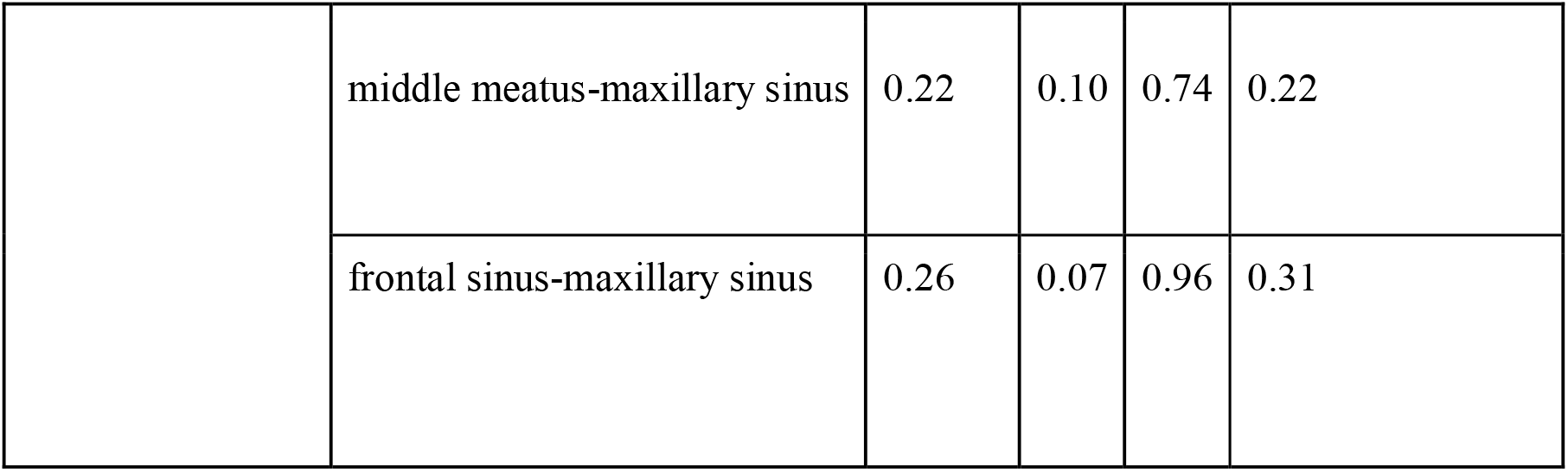
The values of the weighted UniFrac and Bray-Curtis dissimilarity between sampling sites.

We observed statistically significant positive correlations between all intra-individual distances within a patient. The greater disparity between the middle meatus and the maxillary sinus correlated with a greater disparity between the middle meatus and the frontal sinus and between both sinus cavities in the same individual (Bray-Curtis: Spearman’s rho 0.55-0.71, weighted UniFrac: Spearman’s rho 0.58-0.66; p < 0.05).

### The effect of the maxillary ostium size on the microbiome continuity

To assess whether the large middle meatus-maxillary sinus differences noted in some patients were caused by the anatomical separation between the middle meatus and the maxillary sinus, we divided the patients into three groups according to the size of the maxillary ostium. We identified 23 patients with blocked ostium, 14 patients with narrow ostium and 12 patients with wide ostium created during previous surgery.

We found that obstruction of the drainage pathway did not cause larger differences between the microbial communities in the maxillary sinus and the middle meatus. On the contrary, we found that the middle meatus-maxillary sinus distances were smaller in patients with blocked ostia than in patients with narrow or wide ostia. The differences were not statistically significant. However, the difference between the Bray-Curtis metrics in the groups with narrow and blocked ostia was close to the significance cutoff (Mann-Whitney-Wilcoxon test; p = 0.06) with closer middle meatus-maxillary sinus similarity if the ostium was blocked than in case of patent narrow ostium (Fig. 1A).

**Fig. 1.**
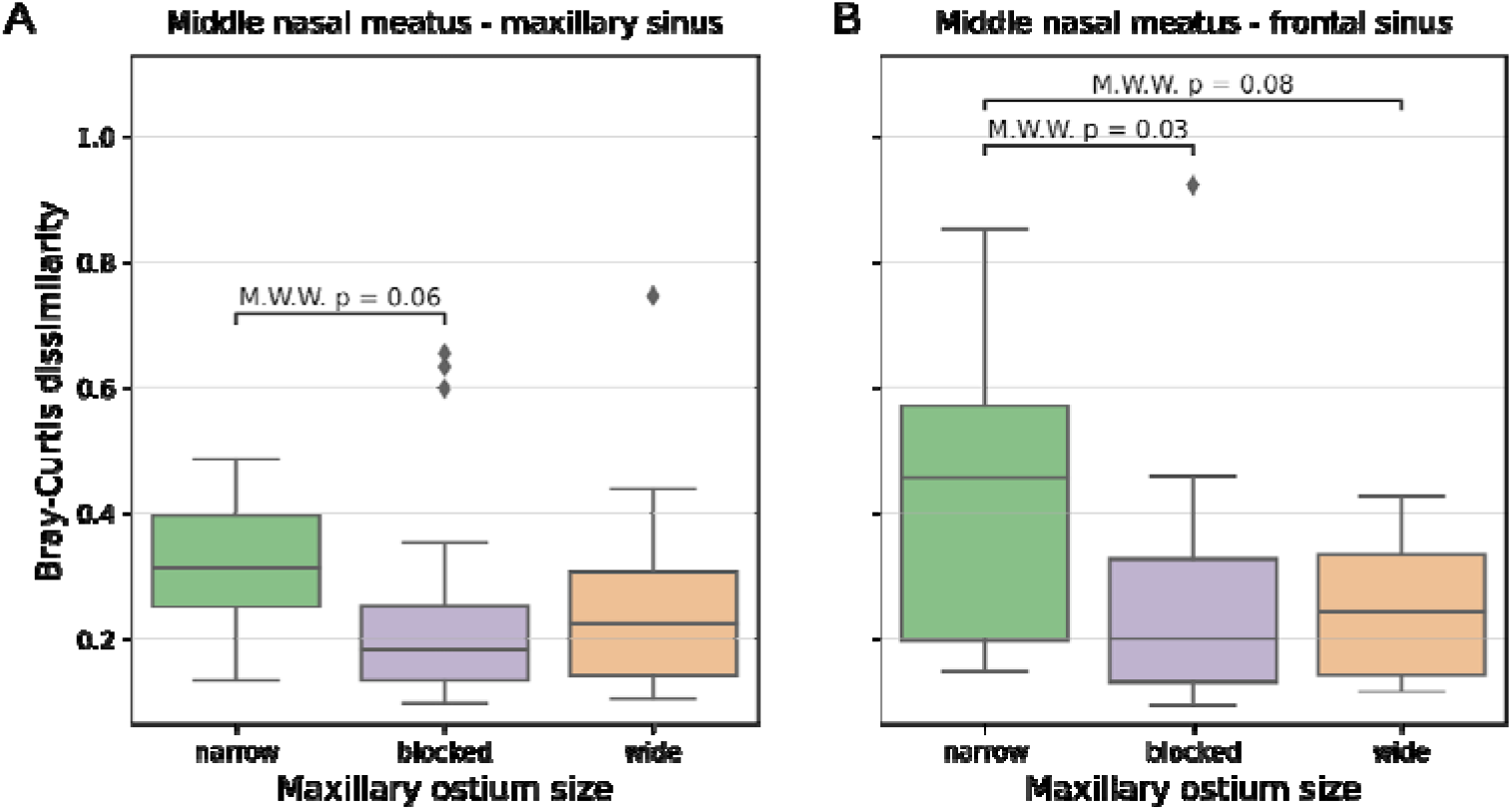
The Bray-Curtis distances between the middle meatus and (A) the maxillary sinus, (B) the frontal sinus in the three subgroups of patients (with wide, narrow and blocked maxillary ostium). Larger metrics values mean greater differences between the sampling sites. M.W.W. - Mann-Whitney-Wilcoxon test.

A similar but stronger relationship with the maxillary ostium size was noted for the middle meatus-frontal sinus beta diversity measures (Fig. 1B). The middle meatus-frontal sinus Bray-Curtis distance in patients with narrow maxillary ostium was significantly greater than in patients with blocked maxillary ostium (Mann-Whitney-Wilcoxon test; p = 0.03) and insignificantly greater than in patients with wide ostium (p = 0.08).

Due to the fact, that only 3 patients in the study group had a patent frontal ostium, analogical computations for frontal ostium did not yield statistically significant results.

### Relationships between intra-individual beta diversity and clinical metadata

Subsequently, we investigated the relationships between the clinical characteristics of the patients and intra-patient beta diversity measures. The distance between the middle meatus and the frontal sinus was significantly smaller in patients with nasal polyps than in patients without nasal polyps (Mann-Whitney-Wilcoxon test; Bray-Curtis distance: p = 0.016, weighted UniFrac distance: p = 0.025). The middle meatus-frontal sinus dissimilarity was also less pronounced in patients with more extensive sinus opacification in the Lund-Mackay score (Spearman’s rho −0.35; p < 0.05). Analogical correlations were not noted for the middle meatus-maxillary sinus distance. Other clinical metadata did not correlate significantly with the beta diversity measures between sampling sites.

### Intra-individual versus inter-individual variability

Despite the high values of beta diversity noted in some patients, PERMANOVA tests (adonis) indicated that in the whole study group the variation between the community structure in different sampling sites was not statistically significant (Bray-Curtis: p = 0.596, weighted UniFrac: p = 0.281). On the contrary, variation caused by differences between patients was significant in both metrics (Bray-Curtis: p = 0.041, weighted UniFrac: p = 0.03). This result indicates that overall the highly pronounced variability between subjects outweighs intra-individual variation of the sinonasal microbiota.

Figure 2 shows that the middle meatus-maxillary sinus and middle meatus-frontal sinus distances within each patient were significantly smaller than distances between the same sites among the study participants. However, the difference between intra-individual maxillary sinus-frontal sinus distances and inter-individual distances was insignificant.

**Fig. 2.**
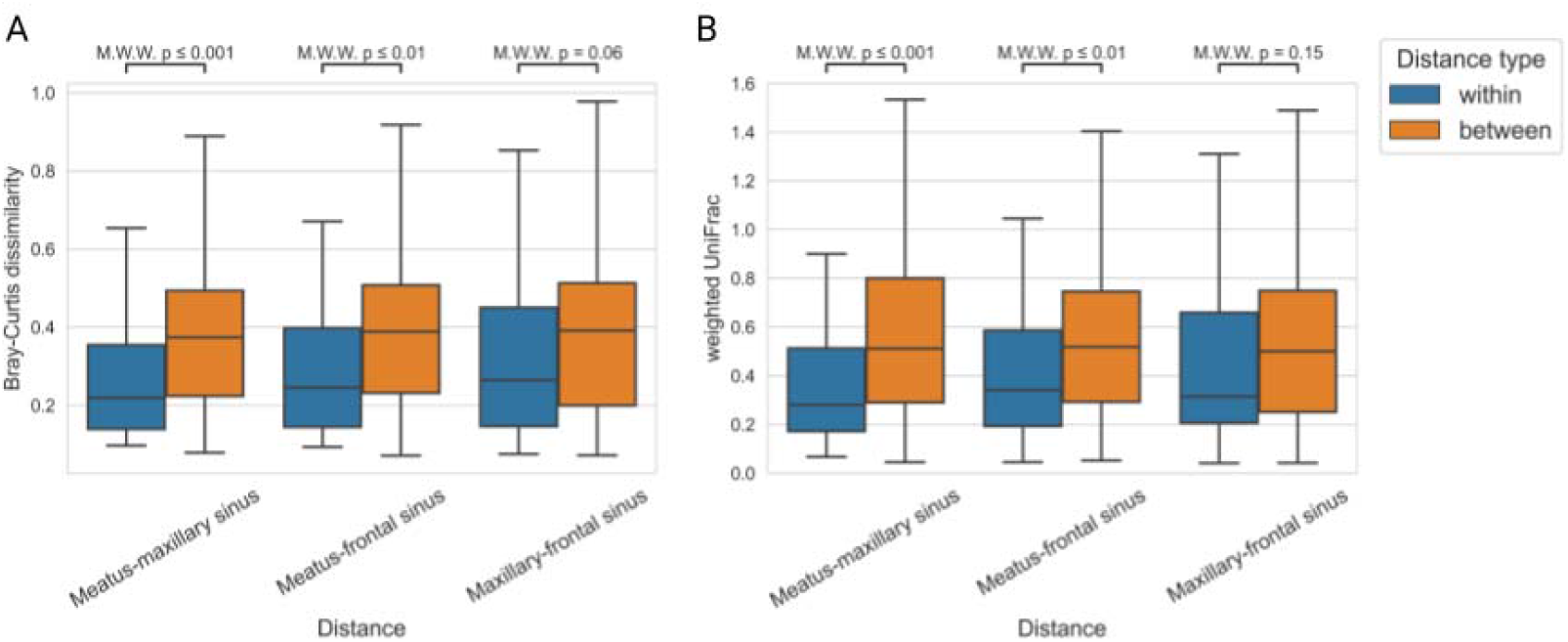
Comparisons of beta-diversity distances within and between patients. The “ within patient” distances were calculated between two sites for each study participant (50 distances). The “ between patient” distances were calculated between one site in each patient and the second site in all other patients (50*49=2450 distances). A. Bray-Curtis distances. B. weighted UniFrac distances. M.W.W. - Mann-Whitney-Wilcoxon test.

In several patients, between-patient Bray-Curtis distances were greater than within-patient distances (Fig. 3). The middle meatus-maxillary sinus differences within a patient were greater than distances between the patient’s middle meatus and the maxillary sinuses of other participants in 7 individuals. The same relationship was observed in 8 patients for middle meatus-frontal sinus distances. A trend towards increased intra-individual variability was observed in patients who also presented higher beta diversity distances from other study participants.

**Fig. 3.**
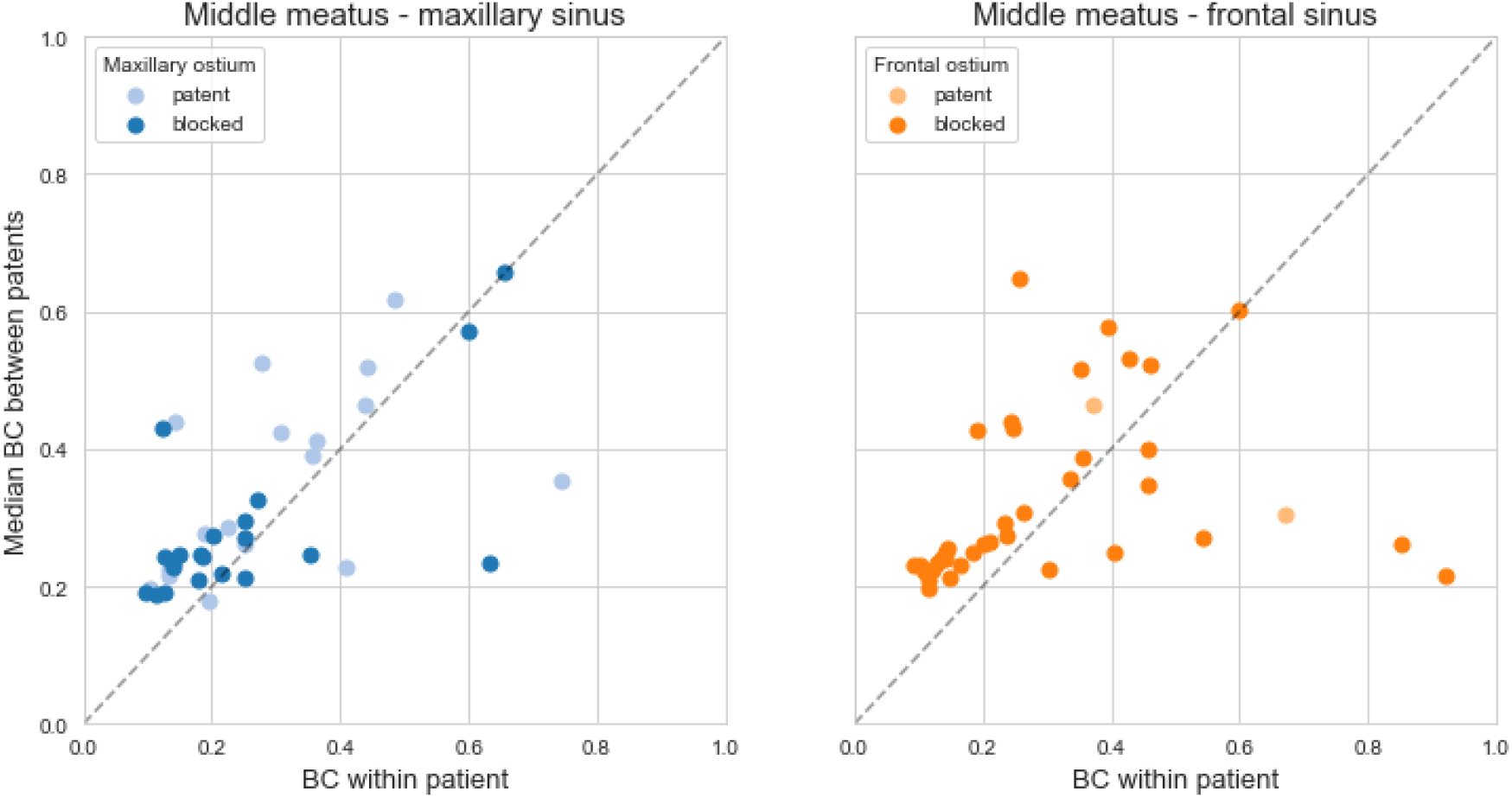
Bray-Curtis beta-diversity distances within and between patients presented for each patient separately. The “ within patient” distances were calculated between two sites for each study participant. The “ between patient” distances were calculated between one site in each patient and the second site in all other patients. The points below the dotted line indicate patients who presented higher variability between sites than between the patient and other study participants.

### Inter-individual functional variability

Phylogenetic Investigation of Communities by Reconstruction of Unobserved States (PICRUSt2) was used to predict the functional potential of the bacterial communities in the samples (18). PERMANOVA analysis of the results revealed statistically significant differences between sampling sites (Bray Curtis dissimilarity based on KO predictions F=2.86, p=0.01; similar values were observed for EC and MetaCyc predictions). The impact of the sinus ostium size on the functional predictions was not significant.

### Alpha diversity in the sinonasal samples

We found no significant differences in alpha diversity (Faith’s phylogenetic diversity) between samples from the middle meatus, the maxillary sinus and the frontal sinus (data not shown). Moreover, there were no differences between alpha diversity in the samples from the maxillary sinuses with wide, narrow and blocked ostia. The taxonomic composition of the samples can be found in the supplementary material (Fig. S1 and Fig. S2).

The relationships between alpha diversity and clinical metadata were not statistically significant. Still, we observed decreased alpha diversity in patients with nasal polyps, a history of recurrent exacerbations or comorbidities such as aspirin-exacerbated respiratory disease, gastroesophageal reflux or asthma. Recent steroid use was almost significantly associated with lower alpha diversity (t-Student test, p = 0.052).

**Fig. S1.**
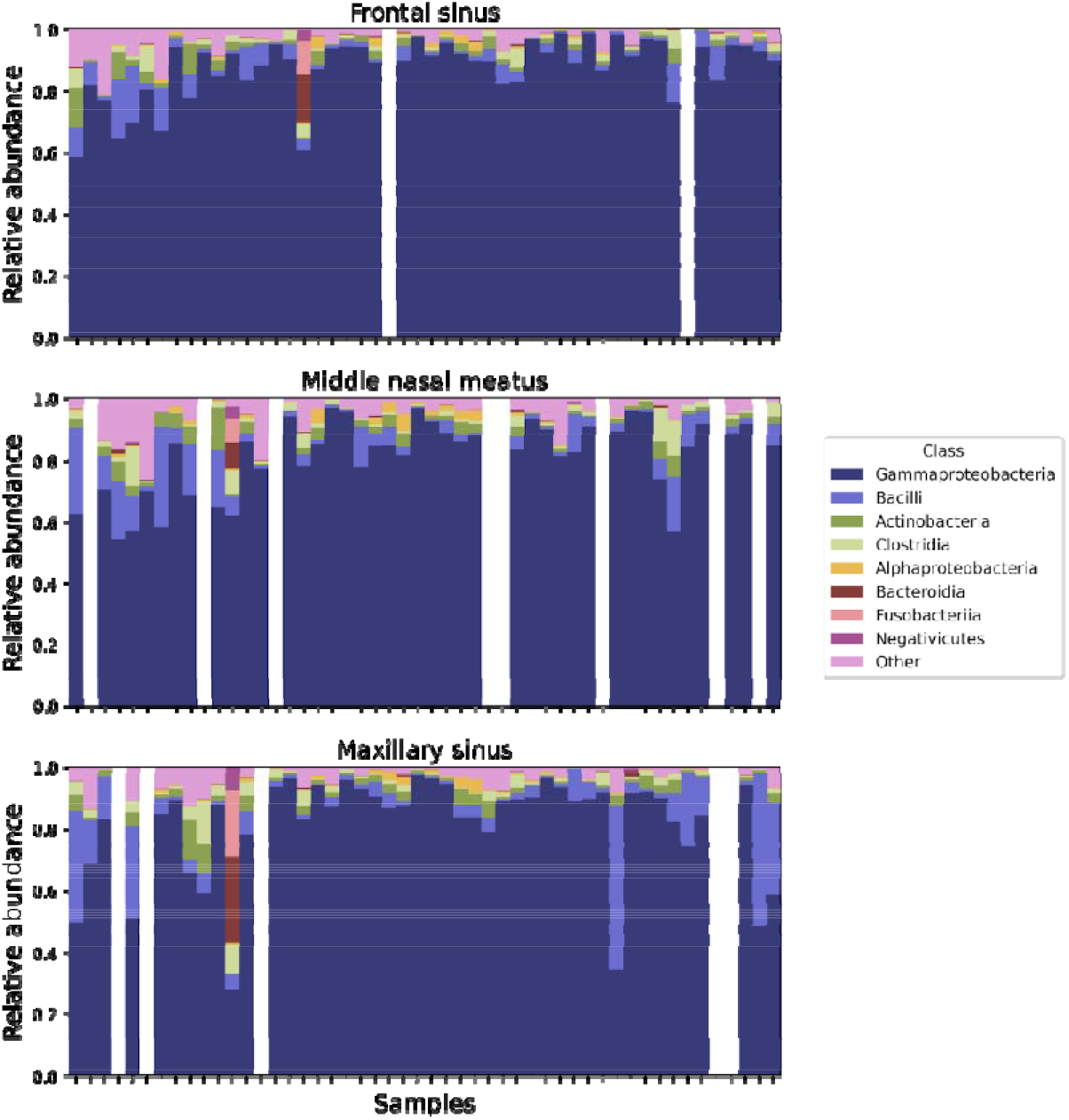
The taxonomic composition of the samples from the frontal sinus, middle nasal meatus and maxillary sinus.

**Fig. S2.**
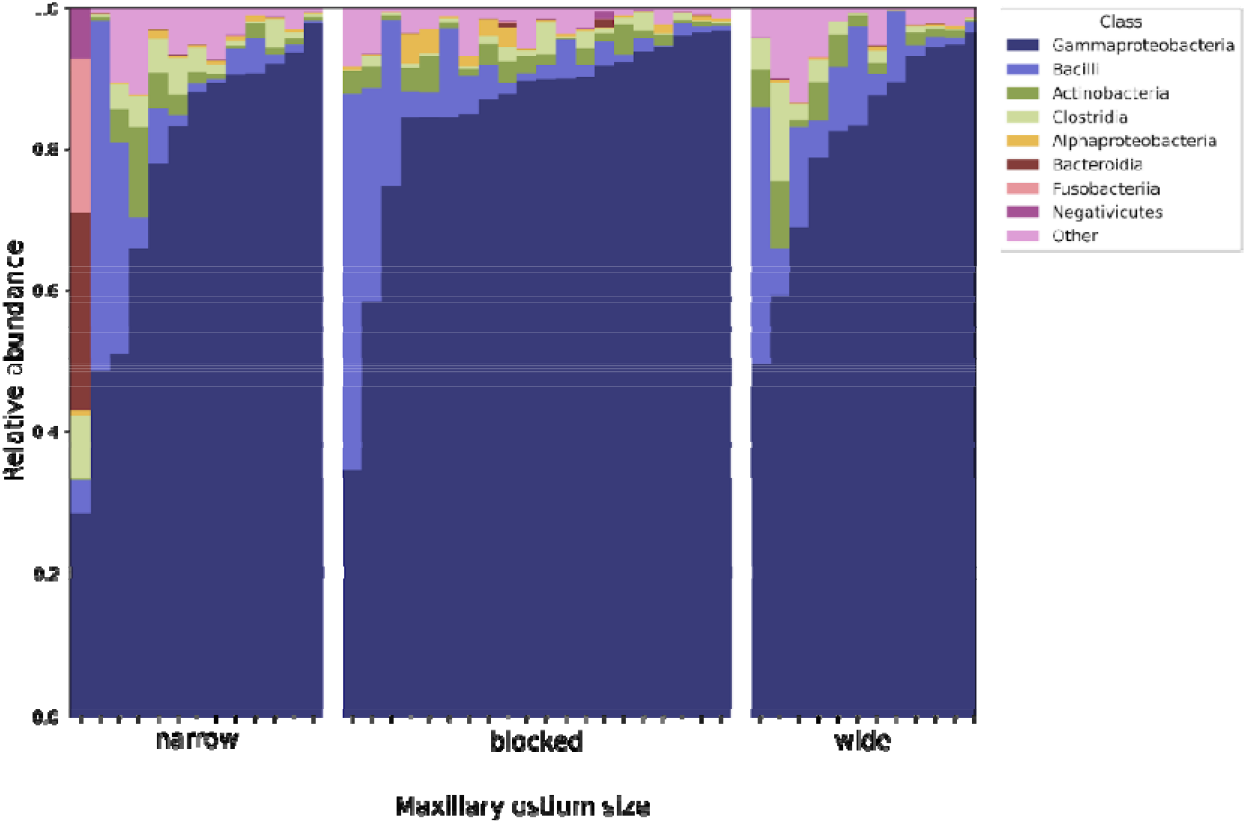
The taxonomic composition of the samples from the maxillary sinus in the groups with wide, narrow and blocked maxillary ostium.

## Discussion

In this study, we explored the topographical differences between the microbial communities in several locations of the sinonasal cavity. We investigated whether the sinonasal cavity could be considered a homogenous habitat or a system of communicating but distinct microenvironments.

Most studies on the subject suggested that the samples from the sinus cavities and the middle nasal meatus were similar (7, 8) and the samples from the middle nasal meatus were representative of the sinuses (1). We decided to question this paradigm because in CRS the drainage into the middle meatus is frequently impaired. It has also been postulated that the hypoxic microenvironment inside an occluded sinus may promote the development of a distinct microbiome while the wide opening of the ostium may decrease the concentration of nitric oxide that controls bacterial growth (7, 19–21). Moreover, our previous study that utilized microbial cultures showed that the middle meatus swabs missed pathogens found in the sinuses in 24% of patients (6). We also found it striking that in several studies of the sinonasal microbiome some patients presented significant differences between sampling sites (7, 11).

These controversies motivated us to explore whether marked heterogeneity within the sinonasal microbiome is frequently encountered and whether it is associated with obstruction of the sinus ostia or driven by other factors. For this purpose, we conducted a detailed analysis of the sequencing data results in the context of the individual patterns of drainage pathway disruption and clinical metadata.

### The microbiome is not universally homogenous across the sinonasal cavity

Differences between samples are quantified by beta diversity measures. However, there is no consensus on what values of these metrics indicate truly meaningful differences. Joss *et al*. assumed that if the weighted UniFrac distance between samples obtained from two sampling sites in the sinuses did not exceed 0.2 then sampling from one site could be considered representative of the other (11). In our study, values of weighted UniFrac above 0.2 were noted in 66% of patients for the middle meatus-maxillary sinus distances and 73% of patients for the middle meatus-frontal sinus distances. Therefore, according to the definition suggested above, the middle meatus samples were not representative of the sinus samples in the majority of patients. The maximal Bray-Curtis value noted in our study reached 0.75 for the middle meatus-maxillary sinus distance and 0.92 for the middle meatus-frontal sinus distance. Joss *et al*. noted between-site weighted UniFrac distances that exceeded 0.2 in 38% of patients while De Boeck *et al*. reported the median Bray-Curtis distance of 0.27 between the maxillary sinus and the ethmoid cells (10). These results show that specimens from a single site may not be representative of the whole sinonasal cavity.

Even though beta diversity metrics are stable within a single experiment and saturate quickly with sequencing depth, they still retain variance dependent upon experiment design (i.e. primer selection) (22), sequencing parameters (i.e. sequencing depth) (23) and data preprocessing pipeline (i.e. denoising method, trimming and rarefaction parameters) (24). The additional layer of complexity is added by a nonlinear nature of common beta diversity measures. All these factors restrict the ability to interpret raw numbers. To alleviate this problem, in our analysis we compare beta diversity between different sampling sites within one patient and beta diversity of the same sampling sites across all patients, utilizing the latter as a point of reference.

### In a minority of patients, the intrapersonal differences outweigh the interpersonal variation

Although the values of beta-diversity measures suggested large between-site dissimilarity in many patients, PERMANOVA analysis did not demonstrate statistically significant differences between sampling sites. The adonis test indicated that interpersonal differences were a significant source of variation, but not the sampling site. There were also no significant differences in alpha diversity between the three locations. The observation that interpersonal variation significantly outweighs intrapersonal variation remains in agreement with the observations of other authors (7, 9-11, 25). Therefore, for certain types of studies, sampling from the middle meatus can be sufficient to represent the composition of the whole sinonasal microbial community (7). De Boeck, who studied a large group of 190 patients with CRS, observed that the sampling site explained a small (2.2%) but statistically significant proportion of microbial variation (10). However, differences were observed between the sinuses and the nasopharynx or the anterior nares. The maxillary sinus and the ethmoid sinus were not significantly different.

Nevertheless, in our study, we showed that in approximately 15% of patients the intrapatient middle meatus-sinus distances were greater than the differences between the patient’s middle meatus and the sinuses of other study participants. (Fig. 3) These observations prove that the apparent intrapersonal variation in some individuals is not captured by statistical analyses that comprise the whole group of CRS patients.

Moreover, PERMANOVA analysis of PICRUSt2 results proved that the functional potential of the bacterial communities differed significantly between various locations within one patient. This observation sheds new light on the sinonasal microbial ecology. It indicates that the microenvironments of the sinuses are heterogeneous and the metabolic activity of bacteria in various niches is distinct. These differences could be explored in detail by deep shotgun sequencing.

### Differences between sampling sites are not related to their anatomical separation

To investigate whether the microbiome heterogeneity was caused by the anatomical separation of the sampling sites, we analysed the configuration of ostia occlusion in each patient (Fig. 1). We expected that blockage of the drainage pathway between the sinus and the middle meatus would result in increased differences between the two sites. Our observations did not support these predictions.

The largest beta diversity distances between the maxillary sinus and the middle meatus were noted in patients with a narrow maxillary ostium. Blocked maxillary ostia were associated with smaller beta diversity between the maxillary sinus and the middle meatus (Fig. 1). Narrow maxillary ostia were also associated with greater distances between the frontal sinus and the middle meatus than blocked or wide maxillary ostia. It suggests that a more divergent sinonasal microbiome was observed in patients whose sinus anatomy was not significantly altered by inflammatory changes (blocked ostia) or by previous surgery (wide ostia). This result was unexpected because other authors reported greater meatus-sinus differences in CRS patients than in healthy individuals and concluded that microbial communities in the sinuses diverge during CRS (9, 25). However, the differences noted in these studies were not statistically significant and require verification in larger study groups.

The results of our study did not support the hypothesis that the microenvironment in an occluded sinus stimulates the development of a distinct microbial community. The analysis of alpha-diversity and PICRUSt2 predictions of the metabolic potential of the microbiome did not reveal any significant microbiome differences between blocked and well-ventilated sinuses. However, the predictions provided by PICRUSt2 are only an approximation of metagenomic functional potential and this phenomenon requires further investigation with the use of deep shotgun sequencing and untargeted metabolomics.

### The degree of sinonasal microbiome variability is characteristic of an individual and not of the topography of the sinuses

Our study showed that between-site beta diversity distances within a patient were positively correlated (patients with greater meatus-maxillary distances also had greater meatus-frontal distances, regardless of the size of the ostia). This observation indicates that a tendency for an increased or decreased between-site variability is a feature that characterizes an individual’s sinonasal microbiome rather than local drainage pathway obstruction between sites.

Patients with nasal polyps or extensive mucosal changes in the sinuses had less dissimilar microbial communities in the middle meatus and the frontal sinus than patients without nasal polyps. Massive sinus opacification and nasal polyps are frequently associated with type 2 inflammation (1). The alpha diversity was not significantly associated with the sampling site or the patency of the sinus ostium. Still, it tended to be lower in patients who presented certain clinical features (recent steroid use, nasal polyps, comorbidities such as asthma, aspirin-exacerbated respiratory disease, and gastroesophageal reflux). Therefore, it is probable that the spatial heterogeneity and composition of the microbiome depend on the individual characteristics of the patient and the type of inflammatory reaction rather than the pattern of ostia occlusion.

## Acknowledgements

TK is supported by the Polish National Agency for Academic Exchange grant PPN/PPO/2018/1/00014.

## Authorship contribution

Joanna Szaleniec – conceptualization, data curation, formal analysis, investigation, methodology, project administration, resources, supervision, visualization, writing – original draft, writing – review & editing

Valentyn Bezshapkin – conceptualization, data curation, formal analysis, investigation, methodology, visualization, writing – original draft, writing – review & editing

Agnieszka Krawczyk - data curation, investigation, methodology

Katarzyna Kopera - data curation, formal analysis, investigation

Barbara Zapala – conceptualization, data curation, formal analysis

Tomasz Gosiewski – conceptualization, funding acquisition, resources, supervision, writing – review & editing

Tomasz Kosciolek – conceptualization, data curation, formal analysis, funding acquisition, investigation, methodology, resources, supervision, writing – original draft, writing – review & editing

## Conflict of interest

The authors declare no conflict of interest.

## Notes

### Competing Interest Statement

The authors have declared no competing interest.

### Summary of Updates

the manuscript has been reorganized and adapted to better fit the needs of a medical audience. no additional results or analyses have been included.

